# bin3C : Exploiting Hi-C sequencing data to accurately resolve metagenome-assembled genomes (MAGs)

**DOI:** 10.1101/388355

**Authors:** Matthew Z. DeMaere, Aaron E. Darling

**Affiliations:** The ithree institute, University of Technology Sydney, 15 Broadway, Ultimo 2007 NSW Australia

**Keywords:** Metagenomics, Hi-C, clustering, next generation sequencing, metagenome-assembled genome

## Abstract

Most microbes inhabiting the planet cannot be easily grown in the lab. Metagenomic techniques provide a means to study these organisms, and recent advances in the field have enabled the resolution of individual genomes from metagenomes, so-called Metagenome Assembled Genomes (MAGs). In addition to expanding the catalog of known microbial diversity, the systematic retrieval of MAGs stands as a tenable divide and conquer reduction of metagenome analysis to the simpler problem of single genome analysis. Many leading approaches to MAG retrieval depend upon time-series or transect data, whose effectiveness is a function of community complexity, target abundance and depth of sequencing. Without the need for time-series data, promising alternative methods are based upon the high-throughput sequencing technique called Hi-C.

The Hi-C technique produces read-pairs which capture in-vivo DNA-DNA proximity interactions (contacts). The physical structure of the community modulates the signal derived from these interactions and a hierarchy of interaction rates exists (*īntra-chromosomal > Inter-chromosomal > Inter-cellular)*.

We describe an unsupervised method that exploits the hierarchical nature of Hi-C interaction rates to resolve MAGs from a single time-point. As a quantitative demonstration, next, we validate the method against the ground truth of a simulated human faecal microbiome. Lastly, we directly compare our method against a recently announced proprietary service ProxiMeta, which also performs MAG retrieval using Hi-C data.

bin3C has been implemented as a simple open-source pipeline and makes use of the unsupervised community detection algorithm Infomap (https://github.com/cerebis/bin3C).

## Background

The number of microbial organisms which can be readily investigated using culture-based techniques is relatively small in proportion to the Earth’s apparent total diversity [1, 2]. Although concerted efforts have found the individual conditions necessary to cultivate a relatively small number of species in the laboratory [3–5], scaling-up this discovery process to the remaining majority is daunting, if not intractable.

Beyond the issue of cultivation, an environmental population can possess at once phenotypic microdiversity and within that group large differences in gene content. With as little as 40% of genes shared within a species [6], this accessory genome is thought to contribute significantly to the dynamics of microbial adaptation in the environment [7–9], Phylogenetic marker surveys (16S amplicon sequencing), while still informative, stand essentially as a proxy for broader discovery processes of the genomic landscape, should they exist. The systematic extraction of entire genomes from an environment will enable a more thorough determination of the constituent species core and accessory gene content (pangenome). The extracted pangenome and community profile will enable investigation of the functional basis of species fitness and niche partitioning within an environment, and further longitudinal experiments will permit studying the dynamics.

Metagenomics offers a direct culture-independent sampling approach as a means to study the unculturable majority. Recent advances in this field have begun to make possible the systematic resolution of genomes from metagenomes; so-called Metagenome Assembled Genomes (MAGs). Tools designed to assess the quality of retrieved MAGs [10, 11] have brought with them suggestions for categorical quality rankings (table 1). Marking an increasing acceptance, the Genomic Standards Consortium (GSC) recently introduced standardised reporting criteria (table 2) for the submission of MAGs to public archives [12], and as of mid-2018 there are more than 5200 MAGs registered in the Genomes Online (GOLD) database [13], As retrieval methodologies improve and new complex environments are studied, the registration rate of new MAGs is expected to eventually exceed that of culture-based studies [12].

**Table 1.** A proposed standard for reporting the quality of retrieved MAGs which uses only estimates of completeness and contamination [10]. Completeness and contamination are independently ranked and are intended to be used in conjunction, e.g. “nearly complete and low contamination”.

| Rank | Completeness (%) | Rank | Contamination (%) |
| --- | --- | --- | --- |
| Near | $\geq 90$ | Low | $\leq 5$ |
| Substantial | $\geq 70$ to $< 90$ | Medium | $> 5$ to $\leq 10$ |
| Moderate | $\geq 50$ to $< 70$ | High | $> 10$ to $\leq 15$ |
| Partial | $< 50$ | Very high | $> 15$ |

**Table 2.** A small component of the reporting details for MAGs as proposed by the Genomic Standards Consortium include ranks of quality [12]. The “finished” rank is left to future advances, while lower ranks are achievable now by Hi-C based genome binning methods. The additional criterion of rRNA genes makes the “high-quality” rank challenging to achieve with current methods.

| Rank | Assembly Quality Criteria |  |
| --- | --- | --- |
| Finished | Single, validated contiguous sequence per replicon without gaps or ambiguities, with consensus error rate or equivalent >Q50. |  |
|  | Completeness and Contamination (%) | Additionally |
| High-quality draft | > 90, < 5 | Presence of 23S, 16S and 5S and $\geq 18$ tRNAs. |
| Medium-quality draft | $\geq 50$, < 10 | |
| Low-quality draft | < 50, < 10 |  |

Most current approaches to the accurate retrieval of MAGs (also called genome binning or clustering) depend on longitudinal or transect data series, operating either directly on WGS sequencing reads (LSA) [14] or on assembly contigs (CONCOCT, GroopM, metaBAT,

MaxBin2, Cocacola) [15–19]. The need for multiple samples can, however, pose a barrier both in terms of cost of sequencing and the logistics of obtaining multiple samples as, for instance, with clinical studies. As an alternative single-sample approach, Hi-C (a high throughput sequencing technique which captures in-vivo DNA-DNA proximity) can provide significant resolving power from a single time-point when combined with conventional shotgun sequencing.

The first step of the Hi-C library preparation protocol is to crosslink proteins bound to DNA *in vivo* using formalin fixation. Next, cells are lysed and the DNA-protein complexes are digested with a restriction enzyme to create free ends in the bound DNA strands. The free ends are then biotin labelled and filled to make blunt ends. Next is the important proximity-ligation step, where blunt ends are ligated under dilute conditions. This situation permits ligation to occur preferentially among DNA strands bound in the same protein complex, that is to say, DNA fragments which were in close proximity *in vivo* at the time of crosslinking. Crosslinking is then reversed, the DNA is purified and a biotin pull-down step employed to enrich for proximity junction containing products. Lastly, an Illumina-compatible paired-end sequencing library is constructed. After sequencing, each end of a proximity-ligation containing read-pair is composed of DNA from two potentially different intra-chromosomal, inter-chromosomal or even inter-cellular loci.

As a high-throughput sequencing adaptation of the original 3C (chromosome conformation capture) protocol, Hi-C was originally conceived as a means to determine, at once, the 3-dimensional structure of the whole human genome [20]. The richness of information captured in Hi-C experiments is such that the technique has subsequently been applied to a wide range of problems in genomics, such as: genome reassembly [21], haplotype reconstruction [22, 23], assembly clustering [24], centromere prediction [25]. The potential of Hi-C (and other 3C methods) as a means to cluster or deconvolute metagenomes into genome bins has been demonstrated on simulated communities [26–28] and real microbiomes [29, 30].

Most recently, commercial Hi-C products ranging from library preparation kits through to analysis services [30, 31] have been announced. These products aim to lessen the experimental challenge in library preparation for non-specialist laboratories, while also raising the quality of data produced. In particular, one recently introduced commercial offering is a proprietary metagenome genome binning service called ProxiMeta, which was demonstrated on a real human gut microbiome, yielding state of the art results [30].

Here we describe a new open software tool bin3C which can retrieve MAGs from metagenomes, by combining conventional metagenome shotgun and Hi-C sequencing data. Using a simulated human faecal microbiome, we externally validate the binning performance of bin3C in terms of adjusted mutual information, and B^3^ Precision and Recall against a ground truth. Finally, for a real microbiome from human faeces, we compare the retrieval performance of bin3C against that published for the ProxiMeta service [30].

## Method

### Simulated Community

To test the performance of our tool on the task of genome binning, we designed a simulated human gut microbiome from 63 high-quality draft or better bacterial genomes randomly chosen from the Genome Taxonomy Database (GTDB) [32]. Candidate genomes were required to possess an isolation source of faeces or feces, while not specifying a host other than human. To include only higher quality drafts, the associated metadata of each was used to impose the following criteria: contig count <= 200, CheckM completeness >98%, MIMAG quality rank of “High” or better and lastly a total gap length < 500 bp. For these metadata based criteria, there were 223 candidate genomes.

In addition to the metadata based criteria, FastANI (v1.0) [33] was used to calculate pairwise average nucleotide identity (ANI) between the 223 candidate genome sequences. As we desired a diversity of species and mostly unambiguous ground truth, a maximum pairwise ANI of 96% was imposed on the final set of genomes. This constraint controlled for the over-representation of some species within the GTDB. Additionally, when two or more genomes have high sequence identity, the assignment process becomes more difficult and error-prone as it challenges both the assembler [34] and creates ambiguity when assigning assembly contigs back to source genomes.

The resulting 63 selected genomes had an ANI range of 74.8% to 95.8% (median: 77.1%) and GC content range of 28.3% to 73.8% (median: 44.1%) (figure 1) (table S1). A long-tailed community abundance profile was modelled using a Generalized Pareto distribution (parameters: shape=20, scale=31, location=0) (figure S2), where there was approximately a 50:1 reduction in abundance from most to least abundant. Lastly, before read simulation, genomes in multiple contigs were converted to a closed circular form by concatenation, thereby simplifying downstream interpretation.

**Figure 1.**
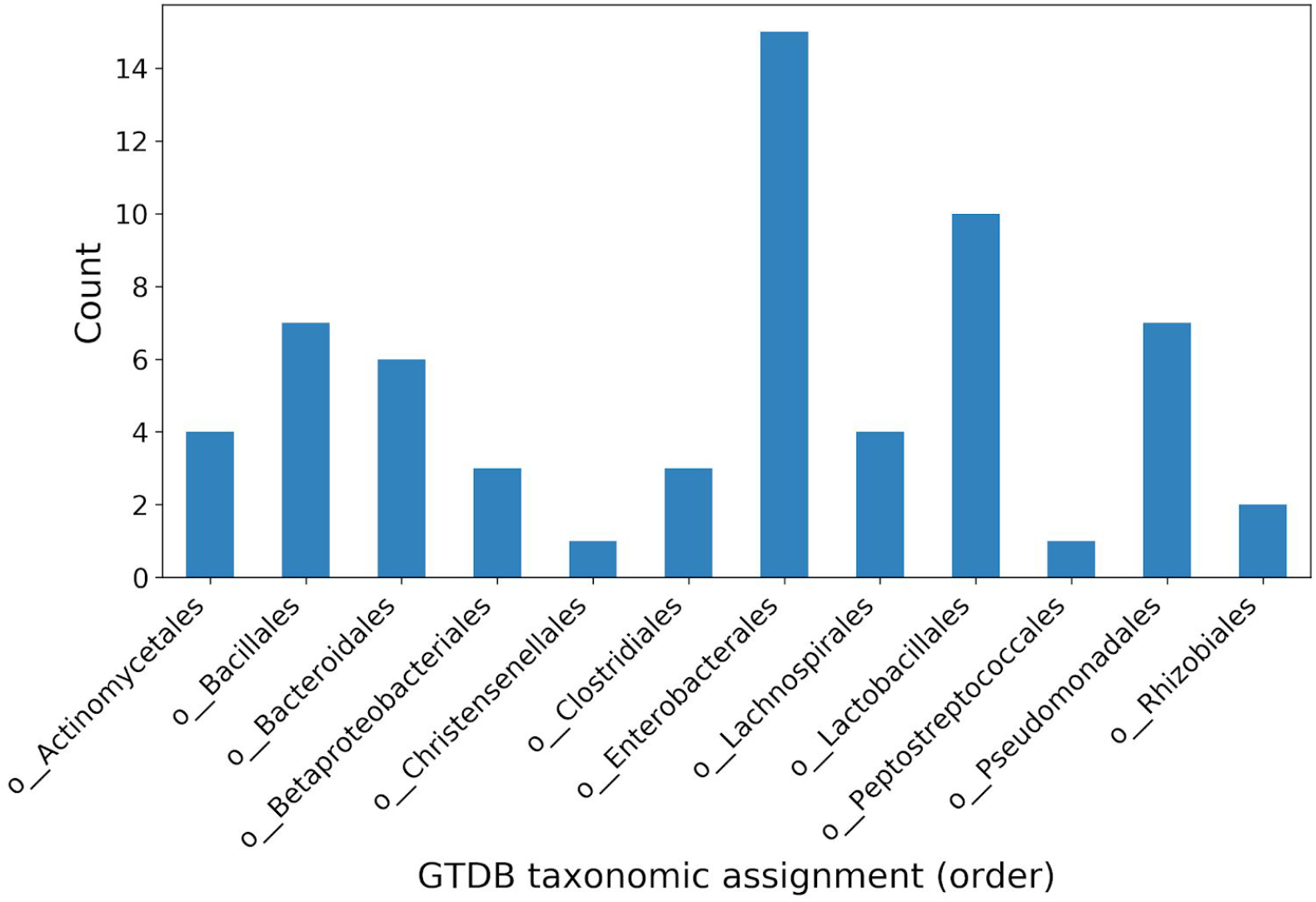
Taxonomic distribution at the order rank of 63 selected bacterial genomes used in the simulated community. The number of each order is a product of the taxonomic distribution of genomes existing in the GTDB, while the constraint that no two genomes be more similar than 96% ANI restricts the over-representation of deeply sequenced species.

### Read-set generation

To explore how increasing depth of coverage affects bin3C’s ability to correctly retrieve MAGs, Hi-C read-sets were generated over a range of depths while keeping shotgun coverage constant. Hi-C depth was parameterised simply by the total number of pairs generated, while shotgun depth was parameterised by the depth of the most abundant community member.

From this definition, an initial read-set with high depth of coverage was produced with 250x shotgun and 200 million Hi-C pairs. The shotgun dataset at this depth constituted 18.2M pairs.

Shotgun reads were generated using the metagenomic shotgun simulator MetaART which wraps the short-read simulator art_illumina (v2.5.1) [35, 36] (options: −M 100 −S 12345 −1 150 −m 350 −s 20 −z 1).

Hi-C reads were generated in two equal parts from two different 4-cutter restriction enzymes (NEB names: MluCI and Sau3AI) using Sim3C [36] (options: −e ${enzyme} −m hic −r 12345 −1 150 --insert-sd 20 --insert-mean 350 --insert-min 150 --linear --simple-reads). Two enzymes were used to mimic the library construction of the real data-set we also analyzed. Repositories containing Sim3C and MetaART can be found at https://github.com/cerebis/sim3C and https://github.com/cerebis/meta-sweeper respectively.

From the initial read-set, a parameter sweep was produced by serially downsampling the initial read-set by factors of 2 using BBTools (v37.25) [37]. The initial Hi-C read-set was reduced 4 times for a total of 5 different depths or 200M, 100M, 50M, 25M, 12.5M pairs (command: reformat.sh sampleseed=12345 samplerate=${d}). In terms of the community genomes, depth of coverage for the subsampling with the greatest reduction factor ranged from 3.5x to 171x for Hi-C.

### Ground Truth Inference

For the task of the whole-community genome binning, a ground truth was constructed by aligning scaffolds resulting from the SPAdes assembly to the “closed” reference genomes using LAST (v941) (Kiełbasa et al. 2011). From the LAST alignments, overlapping source assignment was determined using a methodology we have described previously [34] and implemented as the program alignmentToTruth.py (see availability section). An overlapping (soft) ground truth better reflects the possibility of co-assembly of sufficiently similar regions among reference genomes and the tendency that these regions cause breakpoints in assembly algorithms, leading to highly connected assembly fragments which belong equally well to more than one source.

### Performance Metrics

To validate genome binning, we employed two extrinsic measures; adjusted mutual information (AMI) (sklearn v0.19.2) and weighted Bcubed (B^3^). AMI is a normalized variant of mutual information which corrects for the tendency that the number of agreements between clusters by random chance tends to increase with increasing problem size [38]. Weighted B^3^ is a soft extrinsic metric which, analogous to the F-measure, is the harmonic mean of the B^3^ formulation of Precision and Recall. Here, precision is a measure of cluster homogeneity (like with like), while recall is a measure of the cluster completeness. The B^3^ measure handles overlapping (soft) clusters and better satisfies the constraints that an ideal metric should possess; i.e. homogeneity, completeness, rag-bag and size vs quantity when compared to other metrics. Weighted B^3^ extends the definition to allow the objects under study to have variable values, for which contig length is a natural choice with genome binning problems [34, 39, 40].

In employing two measures, we seek to gain confidence in their agreement while also obtaining the additional insight afforded by the separate facets B^3^ Precision and Recall.

### Real Microbiome

To demonstrate bin3C on real data and make a direct comparison to the proprietary Hi-C based genome binning service (ProxiMeta), we obtained the publicly available high-quality combined whole-metagenome shotgun and Hi-C sequencing data-set used in the previous study [30]. The data-set derives from the microbiome of a human gut (BioProject: PRJNA413092, Acc: SRR6131122, SRR6131123 and SRR6131124).

For this data-set, two separate Hi-C libraries (SRR6131122, SRR6131124) were created using two different 4-cutter restriction enzymes (MluCI and Sau3AI respectively). In using two enzymes, the recognition sites were chosen to be complementary in terms of GC content. When the libraries were subsequently combined during the generation of the contact map, site complementarity provided a higher and more uniform site density over a wider range of target sequence. We conjecture that for metagenome deconvolution, site complementarity is particularly helpful in obtaining a consistent signal from all community members, while higher site density improves recovery of smaller assembly fragments.

All read-sets were obtained from an Illumina HiSeq X Ten at 150 bp. After clean-up (described below), the shotgun read-set (SRR6131123) consisted of 248.8 million paired-end reads, while the two Hi-C libraries consisted of 43.7 million (SRR6131122) and 40.8 million (SRR6131124) paired-end reads.

### Initial Processing

Read clean-up is occasionally overlooked in the pursuit of completing the early stages of genomic analysis. This initial processing step is however essential for optimal shotgun assembly and particularly for Hi-C read mapping where remnants of adapter sequence, PhiX or other contaminants can be a significant noise source.

A standard cleaning procedure was applied to all WGS and Hi-C read-sets using bbduk from the BBTools suite (v37.25) [37]. where each was screened for PhiX and Illumina adapter remnants by reference and by kmer (options: k=23 hdist=1 mink=11 ktrim=r tpe tbo), quality trimmed (options: ftm=5 qtrim=r trimq=10). For Hi-C read-sets, only paired reads are kept to expedite later stages of analysis. Shotgun assemblies for both simulated and read read-sets (table 3) were produced using SPAdes (v.3.11.1) [41] in metagenomic mode with a maximum kmer size of 61 (options: —meta −k 21,33,55,61).

**Table 3.** Assembly statistics for real and simulated human gut microbiomes.

| **Data Set** | **N50** | **L50** | **Contigs**<br>**≥ 1kbp** | **Total**<br>**Contigs** | **Scaffold**<br>**s**<br>**≥ 1kbp** | **Total**<br>**Scaffold**<br>**s** | **Total extent**<br>**(bp)** |
| --- | --- | --- | --- | --- | --- | --- | --- |
| Real human gut | 56,282 | 1277 | 97,760 | 670,379 | 95,521 | 652,723 | 719,550,669 |
| Simulated human gut | 29,009 | 1170 | 24,324 | 116,696 | 23,364 | 41,704 | 240,133,820 |

### Hi-C Read Mapping

As bin3C is not aimed at assembly correction, we opted to use assembly scaffolds rather than contigs as the target for genome binning, electing to trust any groupings of contigs into scaffolds done by SPAdes.

Both simulated and real Hi-C reads were mapped to their respective scaffolds using BWA MEM (v0.7.17-r1188) [42]. During mapping with BWA MEM, read pairing and mate-pair rescue functions were disabled and primary alignments forced to be the alignment with lowest read coordinate (5’ end) (options: −5SP). This latter option is a recent introduction to BWA at the request of the Hi-C bioinformatics community. The resulting BAM files were subsequently processed using samtools (v1.9) [43] to remove unmapped reads, supplementary and secondary alignments (exclude filter: −F 0×904), then sorted by name and merged.

### Contact Map Generation

The large number of contigs (>500,000) typically returned from metagenomic shotgun assemblies for non-trivial communities is a potential algorithmic scaling problem. At the same time, biologically important contigs can be on the order of 1000 bp or smaller, challenging the effective analysis of metagenomic datasets from both sides.

A Hi-C analysis, when conducted in the presence of experimental biases, involves the observation of proximity-ligation events, which in turn rely on the occurrence of restriction sites. The signal we desire to exploit is therefore not smoothly and uniformly distributed between and across all contigs. As a counting experiment, the shortest contigs can be problematic as they tend to possess a weaker signal with higher variance; as a result, they can have a deleterious effect on normalisation and clustering if included. Therefore, bin3C imposes constraints on minimum acceptable length (default: 1000 bp) and minimum acceptable raw signal (default: 5 non-self observations) for contig inclusion. Any contig which fails to meet these criteria is excluded from the clustering analysis.

With this in mind, bin3C constructs a contact map from the Hi-C read-pairs. As in previous work [26], the bins pertain to whole contigs and capture global interactions, which work effectively to cluster a metagenome into genome bins. In doing so, we make the implicit assumption that assembly contigs contain few misassemblies that would confound or otherwise invalidate the process of partitioning a metagenome into genome bins.

bin3C can also optionally construct a contact map binned on windows of genomic extent. These maps are not used in the analysis per se but can be used to plot visual representation of the result in the form of a heatmap (figure S3).

### Bias Removal

The observed interaction counts within raw Hi-C contact maps contain experimental biases, due in part to factors such as mappability of reads, enzyme digestion efficiency, *in vivo* conformational constraints on accessibility, and restriction site density. In order to apply Hi-C data to genome binning, a uniform signal over all DNA molecules would be ideal, free of any bias introduced by the factors mentioned above. Correcting for these biases is an important step in our analysis, which is done using a two-stage process. First, for each enzyme used in library preparation, the number of enzymatic cut sites are tallied for each contig. Next, each pairwise raw Hi-C interaction count *c_ij_* between contigs *i* and *j* is divided by the product of the number of cut sites found for each contig *n_i_, n_j_*. This first correction is then followed by general bistochastic matrix balancing using the Knight-Ruiz algorithm [44].

### Genome binning

After bias removal, the wc-contact map (whole contig) is transformed to a graph where nodes are contigs and edge weights are normalized interaction strength between contigs *i* and *j*. It has been shown that DNA-DNA interactions between loci within a single physical cell (intra-cellular proximity interactions) occur an order of magnitude more frequently than interactions between cells (inter-cellular) [26] and, in practice, the signal from inter-cellular interactions is on par with experimental noise. The wc-graph derived from a microbial metagenome is then of low density (far from fully connected), being composed of tightly interacting groups (highly modular) representing intra-cellular interactions and against a much weaker background of experimental noise. Graphs with these characteristics are particularly well suited to unsupervised cluster analysis, also known as community detection.

Unsupervised clustering of the wc-graph has previously been demonstrated using Markov clustering [26, 45] and the Louvain method [28, 46]. In a thorough investigation using ground truth validation, we previously found neither method to be sufficiently efficacious in general practice [34]. Despite the high signal to noise from recent advances in library preparation methods, accurate and precise clustering of the wc-graph remains a challenge. This is because resolving all of the structural detail (all of the communities) becomes an increasingly fine-grained task as graphs grow in size and number of communities. Clustering algorithms can, in turn, possess a resolution limit if a scale exists below which they cannot recover finer detail. As it happens, modularity-based methods such as Louvain have been identified as possessing such a limit [47]. For Hi-C based microbiome studies, the complexity of the community and the experiment are sufficient to introduce significant structural variance within the wc-graph. A wide variation such aspects as in the size of clusters and weight of intra-cluster edges relative to the whole graph make a complete reconstruction difficult for algorithms with limited resolution.

The state of unsupervised clustering algorithms has however been advancing. Benchmarking standards have made thorough extrinsic validation of new methods commonplace [48], and comparative studies have demonstrated the capability of available methods [49]. Infomap is another clustering algorithm, which like Markov clustering is based upon flow [50, 51]. Rather than considering the connectivity of groups of nodes versus the whole, flow models consider the tendency for random walks to persist in some regions of the graph longer than others.

Considering the dynamics rather than the structure of a graph, flow models can be less susceptible to resolution limits as graph size increases [52]. Additionally, the reasonable time-complexity and the ability to accurately resolve clusters without parameter tuning makes Infomap well suited to a discovery science where unsupervised learning is required.

We have therefore employed Infomap (v0.19.25) to cluster the wc-graph into genome bins (options: −u −z −i link-list −N 10). Genome bins greater than a user-controlled minimum extent (measured in base-pairs) are subsequently written out as multi-FASTA in descending cluster size. A per-bin statistics report is generated detailing bin extent, size, GC content, N50, and read depth statistics. By default, a whole sample contact map plot is produced for qualitative assessment.

In the following analyses, we have imposed a 50 kbp minimum extent on genome bins, partly for the sake of figure clarity and as a practical working limit for prokaryotic MAG retrieval. That is to say, being less than half the minimum length of the shortest known bacterial genome [53], it is unlikely that this threshold would exclude a candidate of moderate or better completeness. If a user is in doubt or has another objective in mind, the constraint can be removed.

## Results

### Simulated Community Analysis

We validated the quality of bin3C solutions as Hi-C depth of coverage was swept from 12.5M to 200M pairs on an assembly (figure 2). A sharp gain in AMI, B^3^ Recall and B^3^ F-score was evident as Hi-C coverage rose from 12.5M to 100M pairs, while the gain between 100M and 200M pairs was less pronounced. Accompanying the upward trend for these first three measures was an inverse but relatively small change in B^3^ Precision. In terms of AMI, the highest scoring solution of 0.848 was at the greatest simulated depth of 200M pairs. Concomitantly this solution had B^3^ Precision, Recall and F-scores of 0.909, 0.839 and 0.873 respectively. For this highest depth sample, 22,279 contigs passed the bin3C filtering criteria and represented 95.4% of all assembly contigs over 1000 bp. There were 62 genome bins with an extent greater than 50 kbp, with total extent 229,473,556 bp. This was 95.6% of the extent of the entire shotgun assembly, which itself was 91.1% of the extent of the set of reference genomes. The remaining small clusters of less than 50 kb extent totalled 1,413,596 bp or 0.6% of the assembly extent (table 3), while unanalyzed contigs below 1000 bp represented 8,103,486 bp or 3.4%.

**Figure 2.**
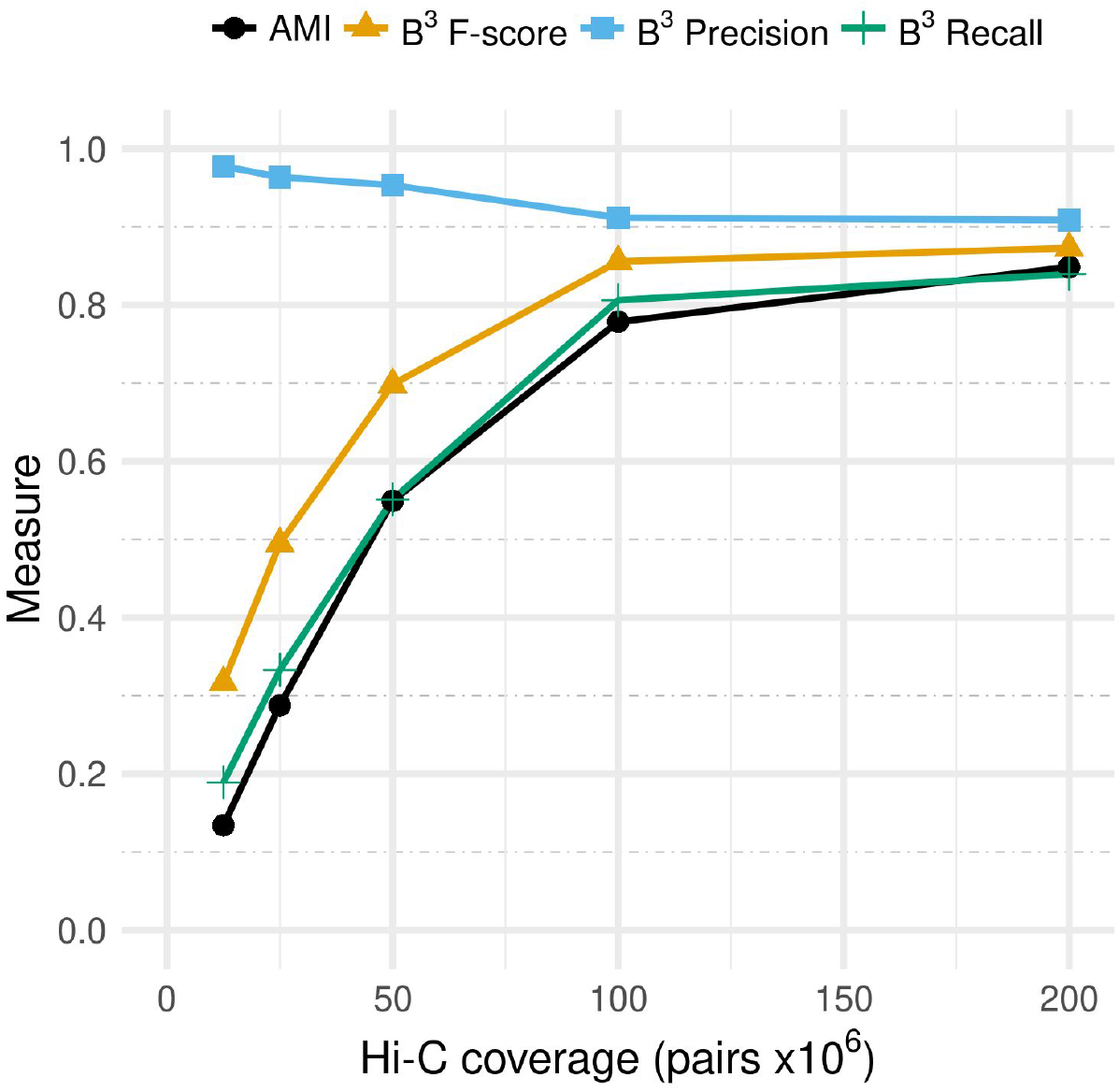
Validation of bin3C solutions using extrinsic measures and a ground truth. bin3C was run against five simulated experiments, with increasing Hi-C depth of coverage while keeping shotgun coverage fixed. With diminishing returns from 100M to 200M pairs, the highest depth of coverage produced the best scoring genome binning solution, with an AMI 0.849 and B^3^ Precision, Recall and F-score of 0.909, 0.839 and 0.873 respectively.

As a soft clustering measure, B^3^ can consider overlaps both within predicted clusters and the ground truth. Regions of shared sequence within our simulated community meant that for 4.4% of assembly contigs the assignment in the ground truth was ambiguous, being shared by two or more source genomes. Meanwhile, bin3C solutions are hard clusters placing contigs in only one genome bin. Even without mistakes, this leaves a small but unbridgeable gap between the ground truth and the best possible bin3C solution. Due to this, when overlap exists in the ground truth, the maximum achievable B^3^ Precision and Recall will be less than unity. Conversely, AMI is a hard clustering measure that requires assigning each of these shared contigs in the ground truth to a single source genome through a coin-toss process. It remains, however, that when bin3C selects a bin for such contigs, either source would be equally valid. For this reason, AMI scores are also unlikely to achieve unity in the presence of overlapping genomes.

Despite these technicalities, a quantitative assessment of overall completeness and contamination is robustly inferred using B^3^ Recall and Precision, as they consider contig assignments for the entirety of the metagenomic assembly. This is in contrast to marker gene based measures of completeness and contamination, where only those contigs containing marker genes contribute to the score. The overall completeness of bin3C solutions, as inferred using B^3^ Recall, rose monotonically from 0.189 to 0.839 as Hi-C depth of coverage was increased from 12.5M to 200M pairs. At the same time, the overall contamination, as inferred using B^3^ Precision, dropped slightly from 0.977 to 0.909. Thus bin3C responded positively to increased depth of Hi-C coverage while maintaining an overall low degree of contamination.

We validated our simulation sweep using the marker gene tool CheckM [10]. CheckM estimated that bin3C retrieved 33 nearly complete MAGs using 12.5M Hi-C pairs, while 39 nearly complete were retrieved using 200M pairs (figure 3). For the deepest run with the most retrieved MAGs, genome bins deemed nearly complete had a total extent which ranged from 1.56 Mbp to 6.97 Mbp, shotgun depth of coverage from 3.34x to 161.2x, N50 from 5797 bp to 2.24 Mbp, GC content from 28.0% to 73.9% and number of contigs from 4 to 787 (figure S4) (table S5).

**Figure 3.**
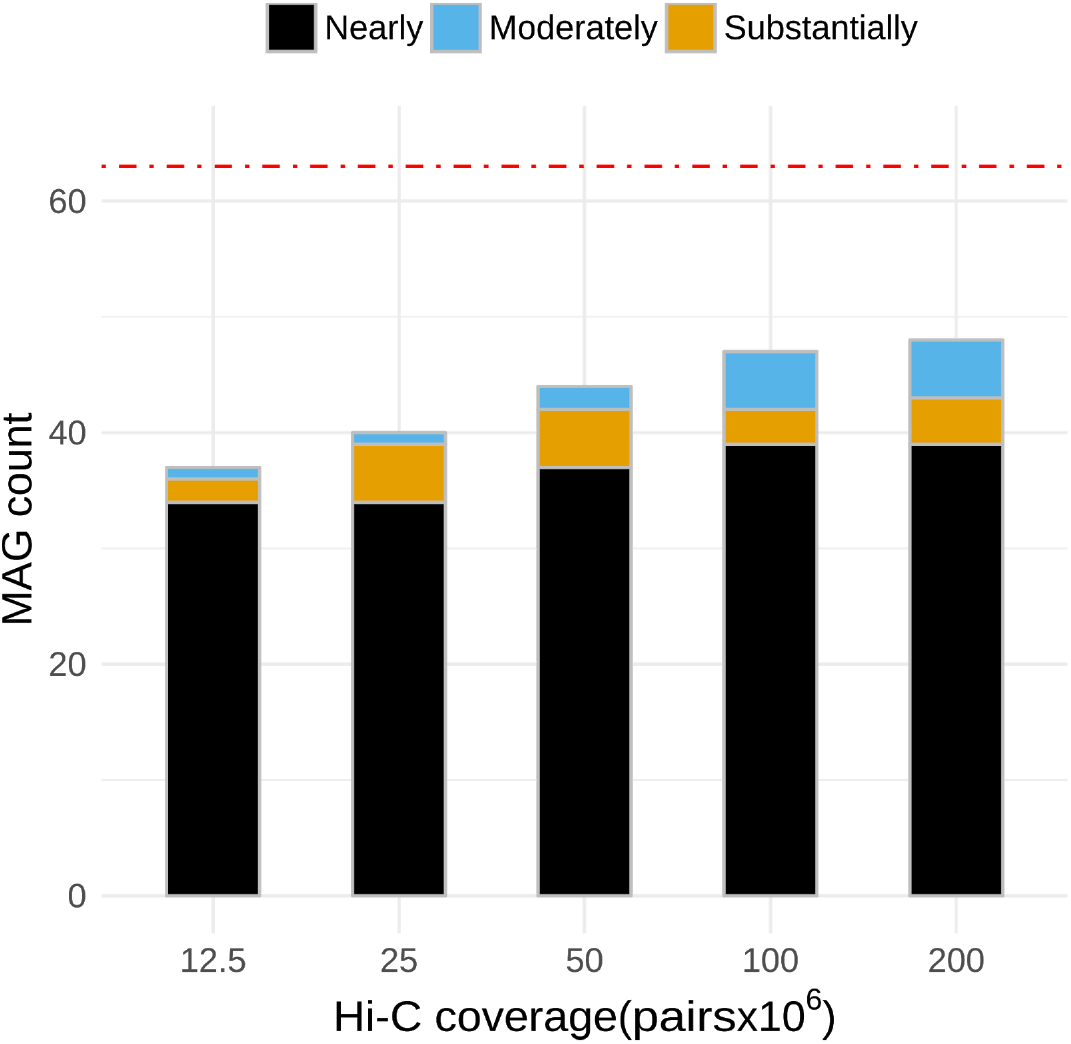
For the simulated community, CheckM was used to validate MAGs retrieved using bin3C for increasing depth of Hi-C coverage. The red dashed line indicates the total number of reference genomes used in constructing the simulated community. The step with the highest depth and consequently highest B^3^ Recall retrieved 39 nearly, 4 substantially and 5 moderately complete MAGs. Nearly complete MAG retrieval at 100M pairs was equal to that of 200M, with 3 substantially and 5 moderately complete MAGs.

Broadening the count to include MAGs of all three ranks: moderate, substantial and nearly (table 1); 37 were retrieved at 12.5M Hi-C pairs, which increased to 48 when using 200M Hi-C pairs. The small increase in the number of retrieved MAGs for the relatively large increase in Hi-C depth of coverage may seem perplexing, particularly in the face of a large change in the extrinsic validation measures AMI, B^3^ Recall and F-score. To explain this, we referred to the cluster reports provided by bin3C, where we found that the average number of contigs in nearly complete MAGs increased from 94 at 12.5M pairs to 179 at 200M pairs. Thus, although marker gene associated contigs are efficiently found at lower Hi-C depth of coverage, obtaining a more complete representation of each MAG can require significantly more depth.

With respect to contamination as inferred by marker genes, CheckM estimated a low median contamination rate of 1.08% across all genome bins with completeness greater than 70%. CheckM, however, also identified four bins where contamination was estimated to be higher than 10% and for which marker gene counting suggested that two genomes had merged into a single bin. We interrogated the ground truth to determine the heritage of these bins and found that each was a composite of two source genomes, whose pairwise ANI values ranged from 93.1% to 95.8%. Each pair shared an average of 131 contigs within the ground truth with an average Jaccard index of 0.19, which was significant when compared against the community-wide average Jaccard of 6.5×l0^−4^. Thus, a few members of the simulated community possessed sufficiently similar or shared sequence to produce co-assembled contigs. Although the co-assembled contigs were short, with a median length of 2011 bp, the degree of overlap within each pair was enough to produce single clusters for sufficiently deep Hi-C coverage. Reference genomes corresponding to two of these merged bins fall within the definition of intraspecies, with pairwise ANI values of 95.80% and 95.85% respectively. The reference genomes involved with remaining two bins are close to this threshold, with ANI values of 93.1% and 93.5%. From this, we would concede that although bin3C is precise, it is not capable of resolving strains.

### Library Recommendations

The time, effort and cost of producing a combined shotgun and Hi-C metagenomic dataset should be rewarded with good results. As bin3C is reliant on both the quality and quantity of data supplied, we felt it important to highlight two factors beyond Hi-C depth of coverage which can influence results.

Shotgun sequencing data forms the basis on which Hi-C associations are made and therefore, the more thoroughly a community is sampled, the better. To demonstrate how this affects bin3C, we reduced the shotgun depth of coverage of our simulated community by half (to 125x) and reassembled the metagenome. Basic assembly statistics for this half-depth assembly were N50 6289 bp and L50 4353. There were 43,712 contigs longer than 1000 bp with an extent of 187,388,993 bp and overall, there were 113,754 contigs with the total extent of 222,522,774 bp. This contrasts to the full-depth (250x) assembly, which had N50 30,402 bp and L50 1105, with 23,364 contigs over 1000 bp with an extent of 232,030,334 bp, and 41,704 total contigs with an extent of 240,133,820 bp. Clearly, the reduction in shotgun depth has resulted in a more fragmented assembly. In particular, the decrease in depth has lead to a 45 Mbp drop in total extent for contigs longer than 1000 bp. This large proportional shift of assembly extent to fragments smaller than 1000 bp is significant as we have found that this length is an effective working limit within bin3C.

We then analysed the resulting contigs with bin3C over the same range of Hi-C depth of coverage as before. Comparison of the AMI validation scores using the half and full depth assemblies (figure 4) shows that, for the more deeply sampled community, bin3C’s reconstruction of the community greatly improved. CheckM estimation of completeness and contamination followed a similar trend (figure S6), where the best result at half depth produced 25 nearly, 4 substantially and 6 moderately complete MAGs, compared against 39 nearly, 4 substantially and 5 moderately complete at full depth.

**Figure 4.**
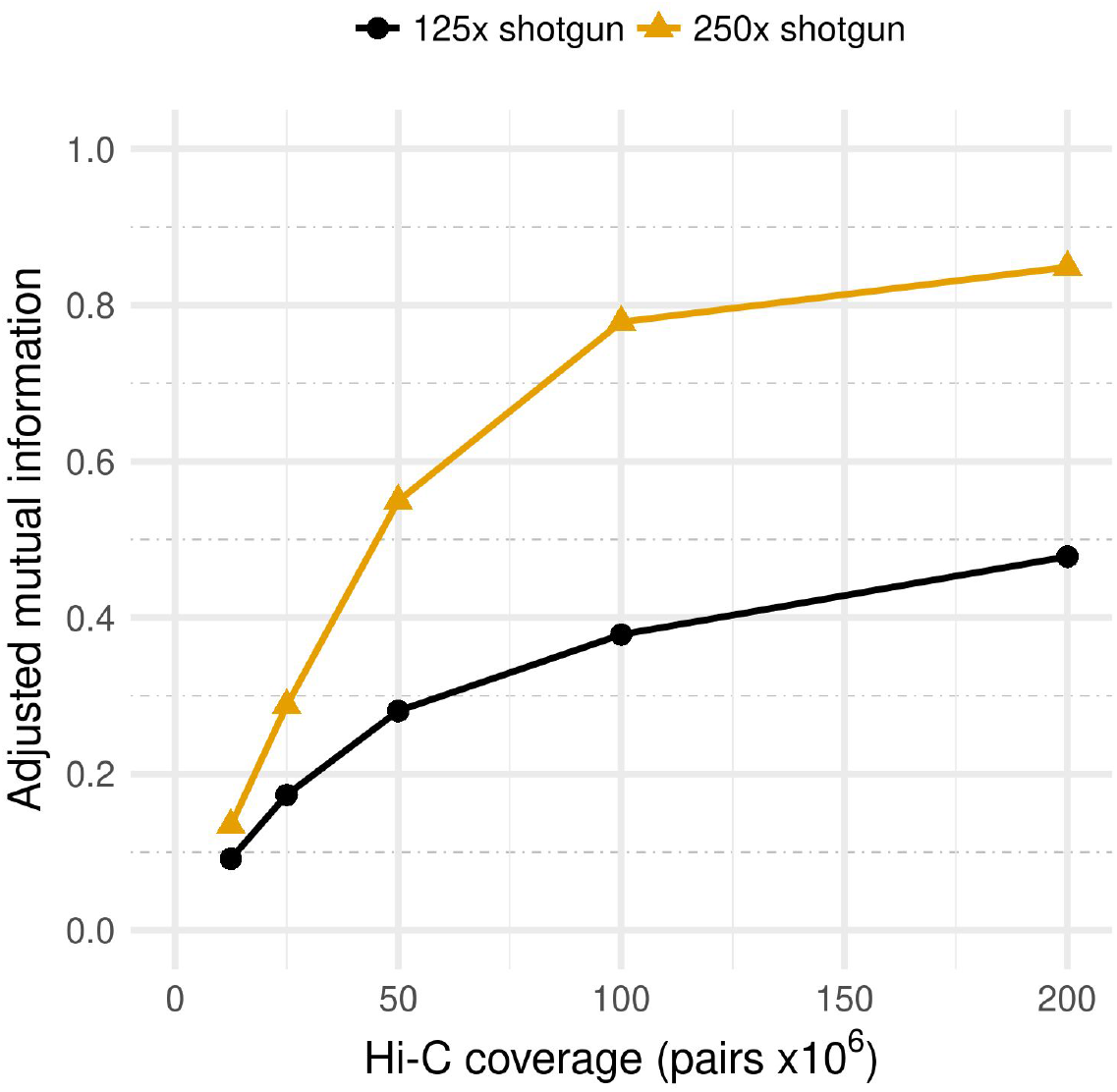
Adjusted mutual information (AMI) scores for bin3C solutions at two different shotgun depths of coverage. For our simulated community, shotgun libraries generated at 125x and 250x coverage demonstrate that although the depth of Hi-C coverage is crucial, so too is the depth of shotgun sequencing.

A recent trend in the preparation of metagenomic Hi-C libraries involves employing two different restriction enzymes during the digestion step [30]. The enzymes are chosen to have different GC biases at their restriction sites. For a microbial community with a diversity of species and consequently a wide range of GC content, the intent of this strategy is more uniform digestion of the extracted DNA, and therefore coverage of Hi-C reads across the metagenome. With wider and more uniform coverage, so the logic goes, should come improved results when performing Hi-C based genome binning.

As our work already involved simulating a two-enzyme library, as used in recent real experiments [30], we elected to repurpose this data to ascertain what gain was had in using two enzymes rather than one alone. The two enzymes used in our simulated libraries are Sau3AI and MluCI. While the Sau3AI restriction site ^GATC is GC balanced, the ^AATT restriction site of MluCI is AT-rich. For our simulated community, source genomes ranged in GC content from 28.3% to 73.8% and their abundances were randomly distributed. For Sau3AI, these extremes of GC content translated to expected cut-site frequencies of 1 in every 338 bp at 28.3% and 1 in every 427 bp at 73.8%. For the less balanced MluCI, the expected cut-site frequencies were instead 1 in every 61 bp at 28.3% and 1 in every 3396 bp at 73.8%. Thus, relative to a naive 4-cutter frequency of 1 in every 256 bp, while the predicted density of sites from Sau3AI is not ideal at either extreme, the site density of MluCI will be very high in the low GC range but very sparse at the high GC range.

For the simulated community full depth assembly, we used bin3C to analyze three Hi-C scenarios: two single enzyme libraries generated using either Sau3AI or MluCI, and a two-enzyme library using Sau3AI and MluCI together. The performance of bin3C was then assessed against the libraries at equal Hi-C depth of coverage using our ground truth. In terms of AMI, the performance of bin3C for the single enzyme libraries was less than that of the combined Sau3 AI+MluCI library (figure 5). Although the gain was small at lower depth, the advantage of a two enzyme model grew as depth increased, where at 100M Hi-C pairs the AMI scores were MluCI: 0.63, Sau3AI: 0.71 and Sau3AI+MluCI: 0.78.

**Figure 5.**
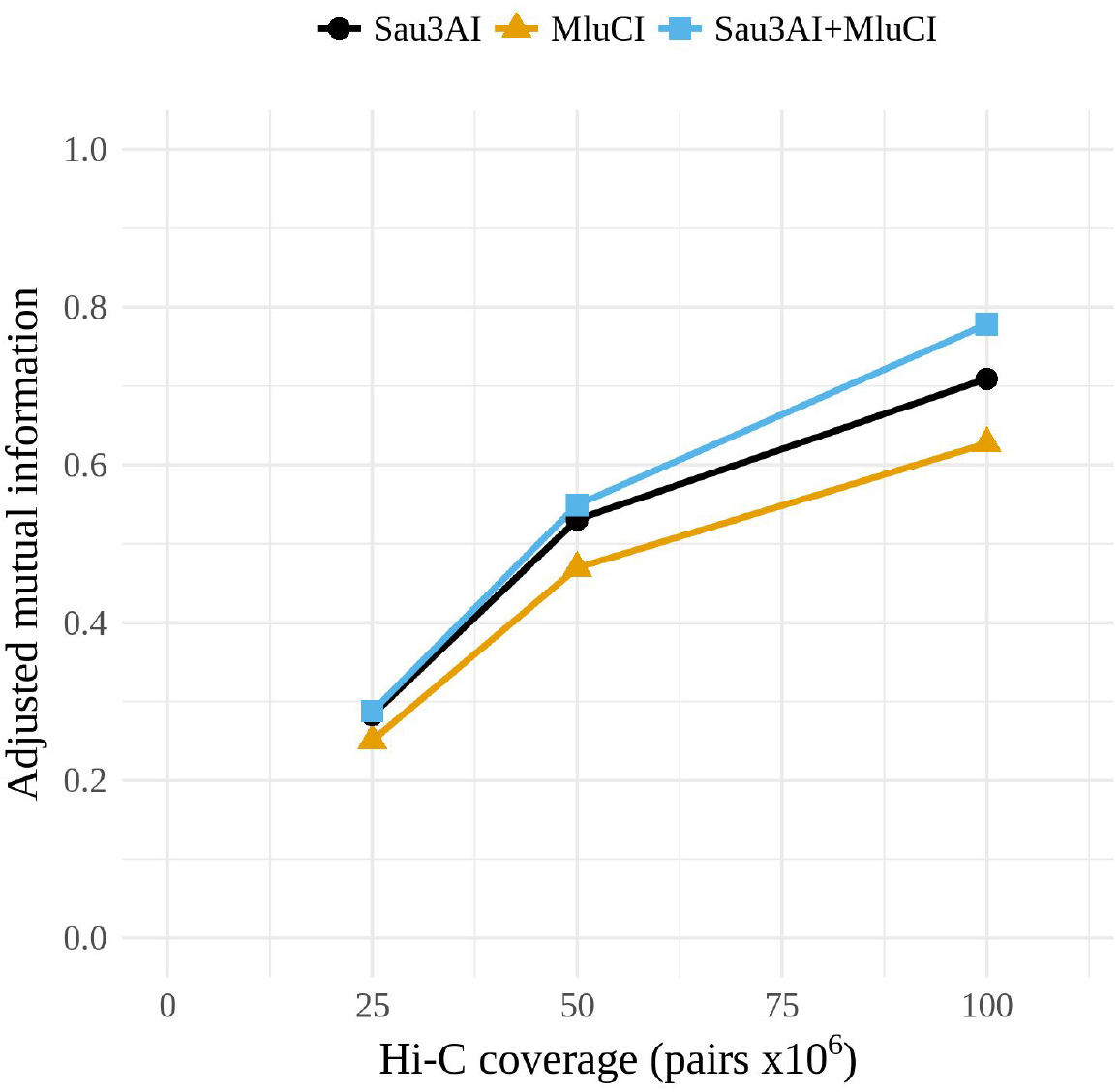
For a simulated community whose GC content varied between 28.3/% to 73.8%, bin3C retrieval performance improved when simulated reads were generated as if from a library prepared using a two enzyme digestion model (Sau3AI+MluCI), rather than if the library was prepared using either enzyme in isolation.

### Real Microbiome Analysis

We analyzed the real human gut microbiome (table 3) with bin3C using the same parameters as with the simulated community along with a randomly generated seed (options: --min-map 60 --min-len 1000 --min-signal 5 −e Sau3AI −e MluCI --seed 9878132). Executed on a 2.6GHz Intel Xeon E5-2697, contact map generation required 586 MB of memory and 15m26s of CPU time, while the clustering stage required 11.6 GB of memory and 9m06s of CPU time. Of the 95,521 contigs longer than 1000 bp, 29,653 had sufficient signal to be included in clustering. The total extent of contigs greater than 1000 bp was 517,309,710 bp for the whole assembly, while those with sufficient Hi-C observations totalled 339,181,288 bp or 65.6% of all those in the assembly.

Clustering the contact map into genome bins, bin3C identified 296 genome bins with extents longer than 50 kbp and 2013 longer than 10 kbp. The 296 clusters longer than 50 kbp had a total extent of 290,643,239 bp, representing 40.4% of the total extent of the assembly, while clusters longer than 10 kbp totalled 324,223,887 bp in extent or 45.1% of the assembly. For clusters greater than 50 kb, shotgun depth of coverage ranged from 3.4x to 498x, N50 ranged from 3119 bp to 297,079 bp, GC content from *28.2%* to 65.0%, total extent from 50,315 bp to 5,460,325 bp and number of contigs from 1 to 495 (table S7).

We analyzed these 296 genome bins using CheckM (figure 6) [10]. For the proposed MAG ranking standard based on only measures of completeness and contamination (table 1), bin3C retrieved 55 nearly, 29 substantially and 12 moderately complete MAGs. In terms of total extent, MAGs ranked as nearly complete ranged from 1.68 Mbp to 4.97 Mbp, while for the substantially complete ranged from 1.56 Mbp to 5.46 Mbp and moderately complete ranged from 1.22 Mbp to 3.40 Mbp (table S8). In terms of shotgun coverage, MAGs ranked as nearly complete ranged from 5.9x to 447.5x, substantially from 4.3x to 416.4x and moderately from 3.7x to 83.4x.

**Figure 6.**
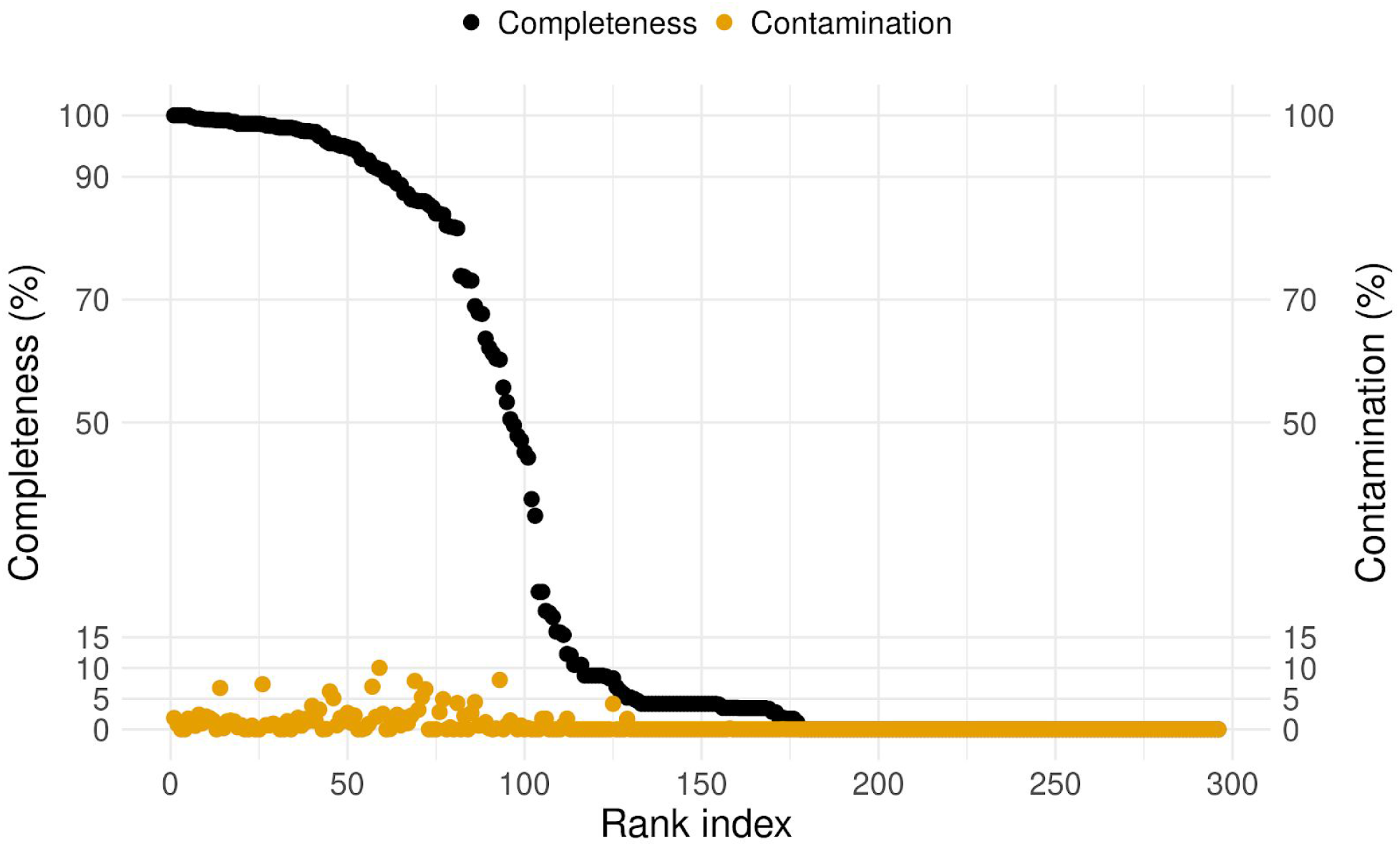
bin3C retrieved MAGs from a real human gut microbiome, ordered by descending estimate of completeness (black circles). Plotted along with completeness is estimated contamination (gold circles). The y-axis grid lines pertain to thresholds used in quality assessment standards: completeness of 50%, 70% and 90% and contamination of 5%, 10% and 15%. Although there is a sharp fall-off in completeness after roughly 75 MAGs, estimated contamination remains consistently low.

Using the more detailed ranking instead from the recently proposed extension to MIxS (table 2) [12], the bin3C solution represented 17 high quality, 78 medium quality and 105 low-quality MAGs. For the high-quality MAGs, shotgun coverage ranged from 10.7x to 447.5x, extent from 1.86 Mbp to 4.10 Mbp (table S9).

### Comparison to previous work

The real microbiome we analyzed with bin3C was first described in a previous study to demonstrate a metagenomic Hi-C analysis service called ProxiMeta [30]. ProxiMeta is the only other complete solution for Hi-C based metagenome deconvolution with which to compare bin3C. As ProxiMeta is a proprietary service rather than open source software, the comparison was made by reanalysis of the same dataset as used in their work (Bioproject: PRJNA413092).

As their study included a comparison to the conventional metagenomic binner MaxBin (v2.2.4) [54], which was one of the best performing MAG retrieval tools evaluated in the first CAMI challenge [55], we have included those results here as well. It should be noted that although MaxBin 2 is capable of multi-sample analysis, all software was run against a single shotgun sequencing sample. We have compared the CheckM validation of bin3C results to the CheckM validation of ProxiMeta and MaxBin as provided in their supplementary data [56],

Regarding the simple ranking standard (table 1), it was reported that ProxiMeta retrieved 35 nearly, 29 substantially and 13 moderately complete MAGs, while MaxBin retrieved 20 nearly, 22 substantially and 17 moderately complete MAGs. On the same metagenomic Hi-C dataset, we found that bin3C retrieved 55 nearly, 29 substantially and 12 moderately complete MAGs (figure 7A). Against MaxBin, bin3C retrieved fewer moderately complete MAGs but otherwise bettered its performance. Against ProxiMeta, bin3C had equivalent performance for the substantially and moderately complete ranks, while retrieving 20 additional nearly complete genomes, representing an improvement of 57%.

In terms of the more complex MIMAG standard (table 2), it was reported that ProxiMeta retrieved 10 high and 65 medium quality MAGs, while MaxBin retrieved 5 high and 44 medium quality MAGs. The bin3C solution retrieved 17 high and 78 medium quality MAGs, which against ProxiMeta represents 70% improvement in high-quality MAG retrieval from the same sample (figure 7B).

**Figure 7.**
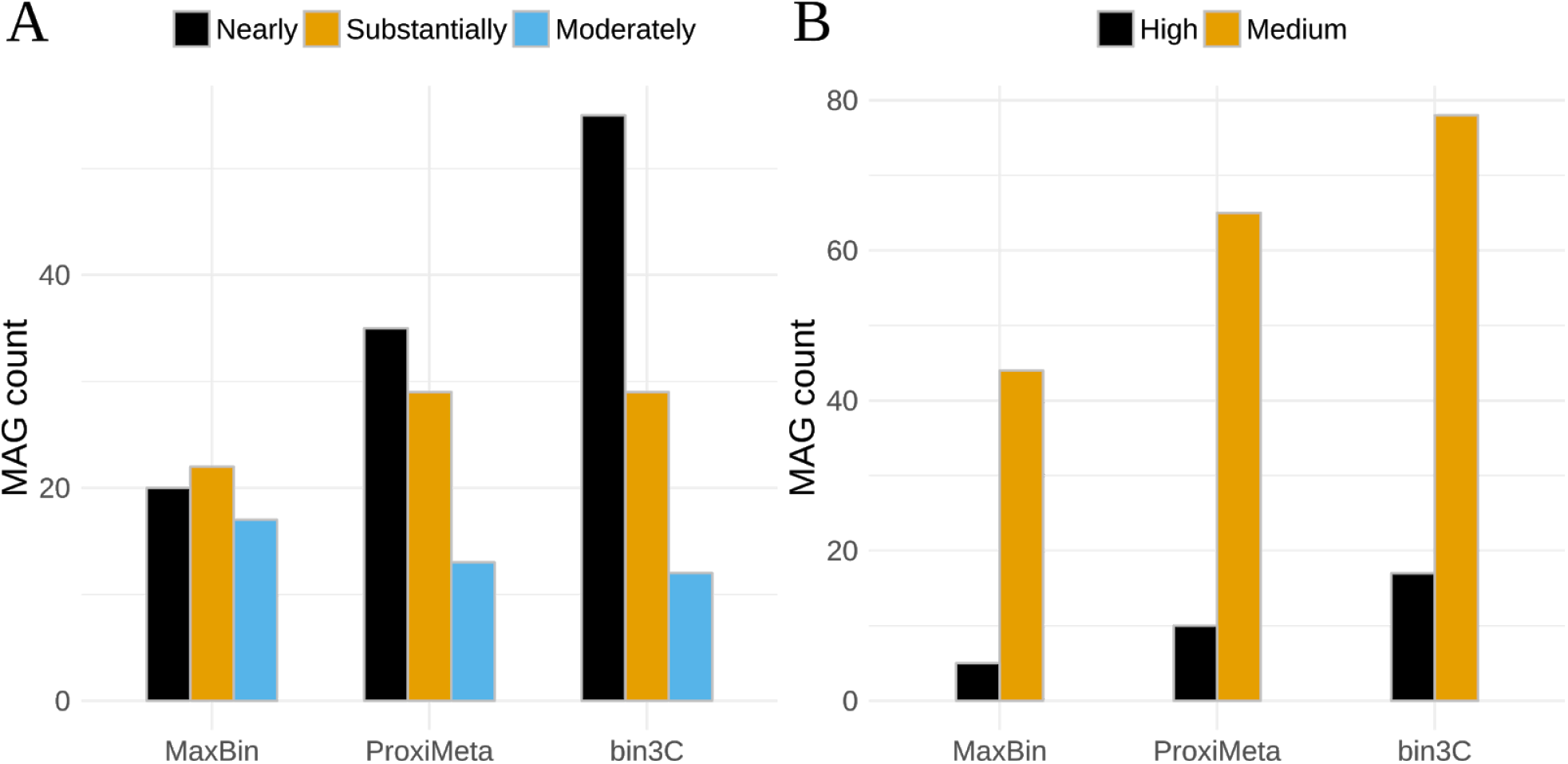
In comparison to existing conventional and Hi-C based single-sample metagenome binning tools, bin3C performs well. When compared by ranking standards, based either on measures of completeness and contamination only (A) [10] or the recent GSC MIMAG reporting standard (B) [12], bin3C retrieves a higher or equivalent number of MAGs in each category. The apparent stringency of the MIMAG high quality is primarily due to the requirement that 5S, 16S and 23 S rRNA genes be present.

It was demonstrated previously that ProxiMeta possessed a higher binning precision than MaxBin and resulted in a much lower rate of contamination [30]. We have found that the precision of bin3C improves on the mark set by ProxiMeta. bin3C’s gains, when retrieving MAGs in the highest quality ranks, are mainly due to the rejection of fewer bins for excessive contamination. For all genome bins over 1 Mbp in extent, bin3C had a median contamination rate of 0.8%, while for ProxiMeta median contamination was 3.5% and MaxBin this was 9.5%.

## Discussion

We have introduced bin3C, an openly implemented and generic algorithm which reproducibly and effectively retrieves MAGs on both simulated and real metagenomic data.

To demonstrate this, we assessed bin3C’s retrieval performance on a simulated human gut microbiome, by way of a ground truth and the extrinsic validation measures of AMI, as well as B^3^ Precision, Recall and F-score (figure 2). bin3C proved to be consistently precise over a wide range of Hi-C depth of coverage, while recall and the overall quality of solutions improved substantially as more Hi-C data was included. Although a high shotgun depth of coverage is not necessary to obtain low contamination MAGs, greater depth of shotgun sequencing has a strongly positive influence on the recall and overall completeness of MAG retrieval (figure 4).

Hi-C MAGs have a characteristically low rate of contamination by foreign genomic content [30]. On a real human gut microbiome, we have shown that bin3C achieves a lower estimated rate of contamination than both the conventional metagenome binner MaxBin [54] and the recently introduced commercial Hi-C analysis service ProxiMeta [30]. For all bins over 1 Mbp as determined by each approach, bin3C’s median contamination rate was 0.8%, while MaxBin was 9.5% and ProxiMeta was 3.5%.

This low contamination rate is a primary reason why bin3C attained the most complete retrieval of MAGs from the real human gut dataset when compared to MaxBin and ProxiMeta (figure 6).

Retrieving 20 more nearly complete MAGs than ProxiMeta, bin3C achieved a gain of 57% on this previous best result (figure 7A). For the stringent GSC MIMAG high-quality ranking, bin3C retrieved 17 MAGs from the gut microbiome, a gain of 70% against the previous best result (figure 7B).

For best results, we recommend that Hi-C metagenomic libraries be constructed using a two enzyme digestion model.

## Limitations and future work

The ground truth as determined in our work is imperfect, notably when a simulated community possesses multiple strains of a single species. The plethora of extrinsic validation measures from which to choose also have their limitations and differences [39, 40, 49]. Though we chose measures which we felt best suited our problem space, these are not in widespread use. Different measures can have significantly different opinions on the agreement between a ground truth and a given solution. Those with the lowest scoring results are not always the most readily chosen for publication.

The use of non-trivial simulated microbial communities makes determining ground truth and measuring accuracy difficult, and yet these are a crucial element of the development process if the resulting methods are to be robust in real experimental use. Under such circumstances, we work from the premise that achieving close to unity on strong validation measures is unlikely to be possible. In our work here, bin3C demonstrated a B^3^ Precision varying between 0.909 and 0. 977, while in work pertaining to metagenome binning with multiple samples, precision values as high as 0.998 were reported using a different formulation of the measure [17]. In practical terms by using CheckM as an operational measure of precision, bin3C achieved a much lower rate of MAG contamination on real data than has previously been reported.

Though marker gene based validation with tools such as CheckM or BUSCO [10, 11] are of great value and easily applied to our work, as validators, their perception is limited only to those sequences which contain marker genes. Ideally, metagenome binning approaches should aim to gather together all the sequence fragments pertaining to a given genome and not only those which contained marker genes. The generalizability of an approach is not assured when the validation measure used in development is systematically insensitive to some aspect of the problem. Therefore, we believe refining the ground truth determination process, to be independent of community complexity, is warranted and would be a useful contribution.

Although bin3C can analyze sequences shorter than 1000 bp, it is our experience that allowing them into the analysis does not lead to improvements in MAG retrieval. We believe the weaker signal and higher variance in the raw observations for Hi-C contacts involving shorter sequences is to blame. A weakness here is relying on the final assembly contigs or scaffolds as the subject of read mapping, where the ends of sequences interrupt alignment. In future work, we believe aligning Hi-C reads to an assembly graph has the potential to achieve better results.

Against the simulated community, the performance of bin3C as indicated by the validation scores AMI and B^3^ Recall, suggests that further gains in retrieval completeness are possible (figure 2).

In particular, strains of the same species can fail to be resolved into separate bins. Improving the resolving power of bin3C or the addition of a *post hoc* reconciliation process to separate these merged bins would be worthwhile.

## List of abbreviations

AMI: adjusted mutual information
ANI: average nucleotide identity
GOLD: Genomes Online Database
GSC: Genomic Standards Consortium
GTDB: Genome Taxonomy Database
MAG: metagenome-assembled genome
MIMAG: Minimum information about a metagenome-assembled genome
MIxS: Minimum information about “some” sequence
3C: chromosome conformation capture

## Declarations

### Availability of data and materials

- Project name: bin3C
- Repository: https://github.com/cerebis/bin3C
- O/S: Linux
- Language: Python 2.7, C/C++
- License: GNU Affero General Public License v3.
- DOI of manuscript version: 10.5281/zenodo.1341423.

Simulators for metagenomic shotgun and Hi-C reads are available at Sim3C repository URL: https://github.com/cerebis/sim3C with the assigned DOI: 10.5281/zenodo.1035049. The Ground truth calculator and shotgun simulator are available at https://github.com/cerebis/meta-sweeper with assigned DOI: 10.5281/zenodo.1341441. Simulated datasets used in this study are available at the assigned DOI: 10.5281/zenodo. 1342169. The real human gut microbiome used in this study was downloaded from the NCBI Sequence Read Archive (http://www.ncbi.nlm.nih.gov/sra) under the accession numbers: shotgun read-set SRR6131123, Hi-C libraries SRR6131122 and SRR6131124 [30]. Supporting material from a previous study used in comparison is available at the assigned DOI: 10.1101/198713.

## Acknowledgements

We thank Professor Steven P. Djordjevic for his gracious support and helpful discussions. This work was supported by the AusGEM initiative, a collaboration between the NSW Department of Primary Industries and the ithree Institute. We acknowledge the use of computing resources from the NeCTAR Research Cloud, the QCIF and the UTS eResearch Group.

## Author contributions

MZD developed the methods, implemented the software, performed and analyzed the experiments, and drafted the manuscript. AED revised and edited the manuscript, and conceived and supervised the project. All authors read and approved the final manuscript.

## Consent for publication

Not applicable

## Ethics approval and consent to participate

Not applicable

